# Evaluation of *Klebsiella pneumoniae* antigen MrkA reveals expression-limited therapeutic potential

**DOI:** 10.1101/2025.08.21.671471

**Authors:** Julia Sanchez-Garrido, Sophia David, Fabio Bagnoli, Monia Bardelli, Carlos Rodrigues dos Reis, David M. Aanensen, Gad Frankel, Joshua L C Wong

## Abstract

Klebsiella pneumoniae (KP) poses a major global health concern, particularly as hospital-acquired strains become increasingly antibiotic resistant. MrkA, the major type 3 fimbrial (T3F) subunit, is a recognised prospective vaccine and monoclonal antibody (mAb) target. We assessed T3F structural (*mrkABCDF*) and regulatory (*mrkHIJ*) gene prevalence and allelic variation across 1649 KP genomes, using 100 diverse isolates for detailed phenotypic characterisation. Despite high conservation MrkA expression itself is variable with genotype failing to predict phenotype. We constructed isogenic KP T3F mutants with absent and supraphysiological MrkA expression to investigate functional consequences and therapeutic responses during infection. In a severe pneumonia model blood dissemination inversely correlated with T3F expression. In vivo antigen profiling revealed only ∼20% of wild-type lung bacteria express MrkA, dropping to ∼5% in blood, with low MrkA expression defining bacteraemic KP irrespective of infection route. Critically, anti-MrkA mAb efficacy was contingent on antigen abundance, with therapeutic benefit restricted to supraphysiological MrkA expression. These findings reveal substantial in vivo MrkA expression variability with critical onward implications for vaccine and mAb development.

## Introduction

*Klebsiella pneumoniae* (KP) is classified as a critical priority pathogen by the World Health Organisation (BPPL, 2024). It is the third most prevalent bloodstream pathogen accounting for 5 to 13% bacteraemias. This proportion increases to 27.9% amongst the sickest patients hospitalised in intensive care units (Holmes *et al*, 2021; Tabah *et al*, 2023). Moreover, KP is the 4^th^ leading global cause of infection-related mortality with 790,000 deaths recorded annually. KP typically infects the lung as the primary site, which can result in sepsis, a life-threatening condition characterised by organ dysfunction resulting from dysregulated host responses (Ikuta *et al*, 2022; Singer *et al*, 2016). KP is typically divided into classical (cKP) and hypervirulent (hvKP) pathotypes, where the former is associated with hospital-acquired infections and antimicrobial resistance (AMR), while the latter, which express unique virulence factors (e.g. siderophores) and thicker capsule, can cause community-acquired infections in healthy individuals (Holmes *et al*., 2021). Notably, hvKP strains expressing AMR genes are on the rise worldwide. Therefore, there is a pressing need to evaluate and develop new treatments strategies for this contemporary threat to human health (Hu *et al*, 2024; WHO, 2024a).

The rise of AMR, with 1.14 million deaths attributable to infections with AMR bacteria in 2021 (of which 12.7% were attributable to KP infection) and the prediction of these numbers increasing to 1.91 million in 2050, illustrates the urgent need to find new therapies beyond small molecule antibiotics (Naghavi *et al*, 2024). Vaccination remains the most effective strategy to combat infections (WHO, 2024b). However, despite the potential of capsule and O-antigen conjugate KP vaccines, most remain in preclinical stages and none are currently approved in humans (Frost *et al*, 2023; WHO, 2024b). Moreover, vaccination requires the induction of a protective immune response that can vary in at risk patient groups and take weeks to months to develop maturely. Given that KP frequently causes disease in hospital settings, often infecting immunocompromised patients and representing the major aetiology in neonatal sepsis, antibody therapy presents itself as an attractive alternative that would provide immediate protection to vulnerable patients (Kumar *et al*, 2023; Stadler *et al*, 2023).

The efficacy and applicability of both vaccination and mAb strategies rely on in vivo expression of the antigens they target. Accordingly, there is a need to establish: i) how widely conserved the antigen is ii) the level of antigen expression at different sites within the host and iii) how accessible the antigen is to antibodies. Here we systematically apply this framework to evaluate MrkA as a prototypical example, using genomic and molecular genetic approaches, in combination with *in vitro* and i*n vivo* profiling and treatment responses.

The implementation and expansion of genomic surveillance has allowed the monitoring of KP lineages across different countries and settings. This includes the determination of AMR and virulence factors together with K (capsule) and O antigen (LPS) type prediction based on sequence data alone (Argimón *et al*, 2021; Lam *et al*, 2022). Whilst KP capsular (K) and/or O-antigens (O) have proven effective as vaccine and mAb targets in preclinical models, their utilisation is limited by the existence of more than 77 K serotypes (147 different loci) and 11 O serotypes (Clarke *et al*, 2018; Lam *et al*., 2022). Thus, there is an increased interest in protein antigens, that may retain species-wide conservation, resulting in broader vaccine and mAb coverage.

Type 3 fimbriae (T3F) are a major KP virulence factor involved in attachment to abiotic surfaces and biofilm formation (Struve *et al*, 2009). T3F are encoded by the *mrk* operon, which includes genes encoding the major fimbrial subunit MrkA, a chaperone (MrkB), an usher (MrkC), the adhesin (MrkD), and a minor fimbrial subunit (MrkF) (Fig. 1A). This structural operon is regulated by the adjacent *mrkHIJ* gene cluster, encoding MrkH, a c-di-GMP-dependent transcriptional activator of *mrkABCDF*, MrkJ, a phosphodiesterase (PDE) that represses MrkH activity, and MrkI, a putative LuxR-type regulator (Johnson *et al*, 2011; Schumacher & Zeng, 2016; Wilksch *et al*, 2011). One promoter lies upstream of *mrkA* (P_mrkA_) driving structural gene transcription. The regulatory genes are controlled by two distinct promoters with P_mrkH_ driving *mrkHI* expression and *mrkJ* expression driven by the cognate P_mrkJ_ promoter (Fig 1A). T3F biosynthesis and regulation is most frequently encoded chromosomally; when plasmid-encoded, T3F operons encode a specific adhesin (MrkD) variant (e.g. KP strain IA565).

**Figure 1.**
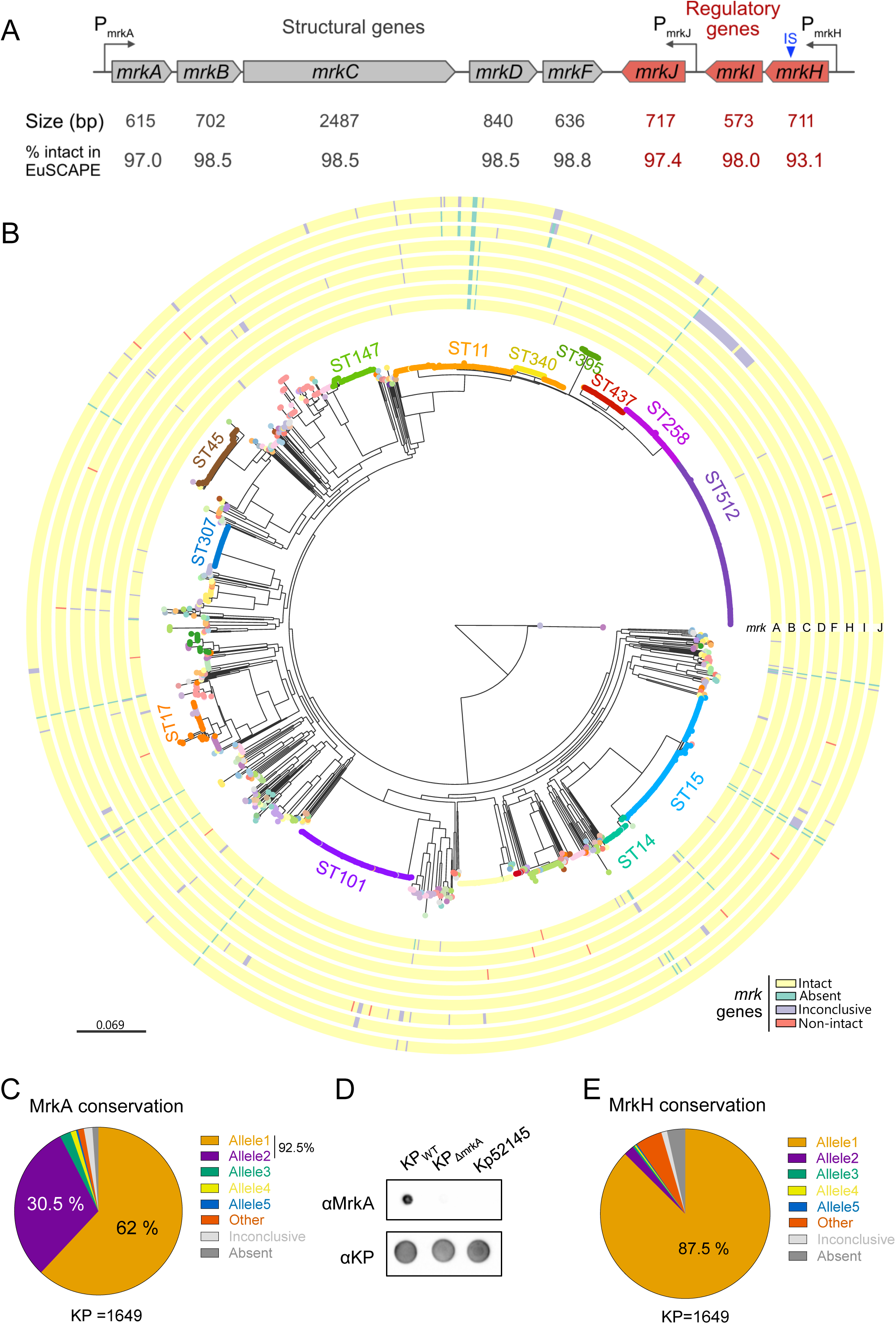
***mrk* genes are very prevalent among circulating KP clinical isolates, with MrkA showing a high degree of conservation** A. The mrk operon consists of structural genes (*mrkABCDF*, grey) and regulatory genes (*mrkH, mrkI, mrkJ,* red). Gene lengths (in base pairs) and the proportion of these genes found intact in the KP isolates of the EuSCAPE collection is shown below each gene. *mrkA*, encoding the major fimbrial subunit, initiates the operon. Regulatory elements (*mrkH*, *mrkI*, and *mrkJ*) modulate expression in response to environmental cues. The position of the insertion sequence (IS) found in Kp52145/B5055 *mrkH* is indicated with an arrowhead (blue). B. Radial phylogenetic tree of 1649 KP isolates with public genomes from the EuSCAPE survey. Isolate tips are coloured by the sequence types (STs) they belong to; significant STs with more than 20 isolates are indicated. Metadata columns from inner to outermost show whether the indicated genes (structural genes *mrkA*, *mrkB*, *mrkC*, *mrkD*, *mrkF* and regulatory genes *mrkH*, *mrkI* and *mrkJ*) are intact (yellow), absent (green), non-intact (orange) or inconclusive (purple). The latter can be caused by multiple hits to the gene (each matching ≥25% of the query gene length) – usually representing fragmentation of the gene within the short-read assembly –, failure to identify the start of the gene (start codon) or the stop codon. The scale bar represents the number of single nucleotide polymorphisms (SNPs) per variable site. A similar interactive visualisation with more detailed metadata is available using Microreact at https://microreact.org/project/mrk-euscape. C. Graph showing the most prevalent MrkA protein sequences in KP from data obtained from the European collection. Allele1 and allele 2 (separated by 1 SNP) account for most of the KP sequences. Inconclusive implies the completeness of the gene could not be determined from the short-read assembly (light grey) and absent (dark grey) denotes no *mrkA*. Details of the 8 most prevalent sequences can be found in Table S2. D. Whole cell samples for the indicated KP strains were spotted and probed with anti-MrkA antibody and anti-KP antibody was used as a loading control. Results confirm that Kp52145 (with truncated mrkH) does not express MrkA, indicating regulatory disruption. The dot blot is representative of n=2 biological repeats. E. Allelic diversity of MrkH among KP genomes. Pie chart shows that 87.5% of KP genomes encode the same MrkH protein variant (allele 1). Inconclusive implies the completeness of the gene could not be determined from the short-read assembly (light grey) and absent (dark grey) denotes no *mrkH*. Details of the 8 most prevalent sequences can be found in Table S2.

Previous genomic studies have reported a high prevalence of the T3F in *K. pneumoniae*; however, these analyses were limited by small genome collections (typically 50–300 isolates) and often focused solely on the presence of structural genes (Bialek-Davenet *et al*, 2014; Onori *et al*, 2015) or individual components such as *mrkA* and/or *mrkD* (Alcántar-Curiel *et al*, 2013; Vuotto *et al*, 2017). The identification of anti-MrkA antibodies in the sera of humans and mice previously exposed to KP has led to MrkA being considered as a putative vaccine and mAb candidate. Indeed, both preclinical MrkA vaccines and mAbs have been shown to be effective in protecting mice against high-dose pneumonia challenge with various KP isolates (Berry *et al*, 2024; Wang *et al*, 2016; Wang *et al*, 2017). These results, coupled with the polymerisation of the MrkA pilin subunit within the T3F (i.e. representing high numbers of binding sites per pilus), the conservation of MrkA in chromosomal and plasmid contexts (Wang *et al*., 2016) and the role of T3F in biofilm formation (Chu *et al*, 2023) make MrkA an attractive target for therapeutic applications. In order to realise this potential, a better understanding of its role in KP pathogenesis is required, which includes defining MrkA expression levels at both the primary site of infection and following dissemination. In particular, as KP is a leading cause of bacteraemia, evaluating any potential therapeutic antigens expression in the blood stream is key.

We found that both structural and regulatory *mrk* operons are highly prevalent and conserved in most contemporary clinical KP isolates. However, the high-risk cKP clones ST101 and ST258 were found to exhibit more variability. We then bridge the gap between genomic presence and phenotypic MrkA expression by multimodally profiling of a species-wide collection of representative clinical isolates. Whilst this combined approach confirmed high levels of T3F gene conservation it also highlighted strain-to-strain variation in MrkA expression, a feature that was independent of capsule-type and mucoviscosity. By generating KP mutants lacking (KP *_mrkA_*) or over expressing (KP *mrkJ* premature stop codon – *mrkJ\**) MrkA we show that MrkA is not required for establishing infection in the lungs or for the spread to the blood in acute mouse KP infection models. We found that while ∼20% KP express T3F in the lung, this drops to only ∼5% in the blood, with low expression also encountered in the liver and spleen. Finally, we show that the efficacy of anti-MrkA mAb therapy is dependent on MrkA expression levels as mice infected with KP *mrkJ\** were better protected. These results suggest that selecting antigens for vaccination or mAb therapeutics cannot rely on genomics alone and that functional in vitro and in vivo validation is also needed. Moreover, our results suggest the mAbs against MrkA may have limited benefit and should only be employed in the context of combination therapy.

## Results

### MrkA and the Structural Operon

To evaluate the potential of the type 3 fimbriae (T3F) subunit MrkA as a therapeutic target in *Klebsiella pneumoniae*, we first assessed the prevalence of the structural *mrkABCDF* operon in a collection of 1717 short-read assembled genomes from the KP species complex. These clinical isolates were collected in hospitals across Europe in 2013-14 as part of the European Survey of Carbapenemase-producing Enterobacteriaceae (EuSCAPE) (David *et al*, 2019). Our analysis focused on *K. pneumoniae sensu stricto* (KP), which comprised the vast majority of genomes (1649/1717). Using a BLASTn approach (Camacho *et al*, 2009) (see *Methods*), we found that all five *mrkABCDF* genes were intact in 95.8% (1579/1649) of KP genomes (https://microreact.org/project/mrk-euscape) (Fig. S1A). The most commonly disrupted gene was *mrkA*, which was either absent or truncated in 22 isolates. These, along with other isolates lacking structural *mrk* genes, formed singletons or small clusters in the phylogenetic tree, suggesting that loss of T3F may limit clonal expansion (Fig. 1B).

We identified 17 protein level allelic variants of the MrkA (Fig. 1C; Table S1, S2). The dominant variant, Allele 1, was found in 62.0% of genomes (1022/1649) and is also encoded by our prototype KP strain ICC8001 (KP_WT_), an ATCC43816 derivative (Wong *et al*, 2019) which has been previously shown to express MrkA (Willsey *et al*, 2018). Allele 2 differed by a single non-synonymous mutation (T28N). Together, these two alleles accounted for 92.5% of isolates, indicating limited protein sequence diversity (Fig. 1C; Table S1, S2). Genetic analysis of the immediate *mrkABCDF* upstream region revealed no variation in key motifs including the ribosome-binding site (RBS) and canonical-10 and-35 promoter elements in 98.6% of genomes in this large collection (Wilksch *et al*., 2011). However, a small subset of 23 isolates harboured deletions directly upstream of *mrkA*, distributed across multiple lineages including ST11, ST258/512, ST101, and ST15 (Table S1, Fig. S1B).

### MrkH and the Regulatory Gene Cluster

We next examined the regulatory *mrkHIJ* gene cluster, which modulates T3F expression via the transcriptional activator MrkH (Fig. 1A). We aimed to provide a comprehensive *mrkHIJ* analysis as this has been largely overlooked in surveillance reporting and these gene products are predicted to be mandatory for T3F expression and regulation in KP. The three regulatory genes were identified in 92.8% (1531/1649) of genomes, with 98.4% of these (1507/1531) encoding intact open reading frames (Fig. 1B; Fig S1A, Table S1). The most commonly undetected gene was *mrkH* (93/1649 genomes). Interestingly, we also observed a disruption of *mrkH* in the lab strain Kp52145 (Calderon-Gonzalez *et al*, 2024; Lery *et al*, 2014) where a 1.34 kb insertion sequence (IS3) element interrupts *mrkH* at nucleotide position 525. The structural operon and other regulatory genes remain intact in this isolate. We evaluated Kp52145 T3F expression by dot blot, probed with a polyclonal MrkA antibody, and employed KP_WT_ and KP_Δ*mrkA*_ – which lacks the structural *mrkA* gene – as positive and negative controls respectively. Absent MrkA expression was found in Kp52145 indicating that MrkH is essential for T3F expression (Fig. 1D). On closer inspection of the 93 genomes with *mrkH* unconfirmed or absent *mrkH*, 70 had multiple partial matches to *mrkH* – each covering >25% of the query sequence length (Table S1). This is highly suggestive gene fragmentation potentially driven by IS insertion as observed in Kp52145. Indeed, an insertion sequence (IS) was identified as the cause of the assembly disruption in *mrkH* in a single clade of ST258 encompassing 67.1% of these genomes (47/70) (Fig 1B, Fig. S1D). Our data indicate that, despite an otherwise full *mrk* structural and regulatory gene complement, these isolates will not express T3F. Nonetheless, the expansion of this clade suggests that IS-mediated disruption of *mrkH* does not carry a strong fitness cost within this lineage.

For completeness we then proceeded to assess protein level variability in MrkH. Whilst we identified 45 different MrkH alleles, one (Allele 1) was highly dominant, present in 87.5% of isolates (Fig. 1E; Table S2). This suggests that while some variation exists, most strains encode a highly conserved form of MrkH. Upstream genetic sequence analysis of the *mrkHI* operon showed no variation in the RBS or the-10 and-35 promoter elements (Tan *et al*, 2015) and a high level of conservation (Fig. S1F). However, the *mrkHI* promoter was fragmented at the same breakage site in a clade of 35 ST101 isolates. (Fig. S1G). Long-read sequencing of one representative isolate confirmed IS-mediated disruption of the promoter. These 35 genomes represent 28.2% (35/124) of the ST101 lineage, which is a clinically significant clonal group rising in prevalence across Europe (Budia-Silva *et al*, 2024; Campos-Madueno *et al*, 2022) (Fig. S1G) and may exhibit limited T3F expression due to transcriptional *mrkH* suppression.

In summary, our high-level genomic evaluation demonstrates that 88.4 % of strains encode intact structural (m*rkABCDF*) and regulatory (*mrkHIJ*) genes. The most commonly disrupted structural gene is *mrkA* and the most commonly disrupted regulatory gene in *mrkH*. There is very little protein level variation in MrkA and MrkH and presence of both is required for T3F expression in KP.

### Phenotypic characterisation of MrkA expression

In order to investigate the role of MrkA and define its expression during infection, we next used isogenic mutants we generated in our KP_WT_ strain: KP_ΔmrkA_ and KP_mrkJ*_, in which a premature stop codon was inserted at position 10 (Ile10stop) of the negative regulator *mrkJ* (Fig. 2A, Fig. S2A). We validated the T3F mutant phenotypes by dot-blot of whole bacteria grown in M9 minimal media with 0.4% glucose (M9-Glc) (Schurtz *et al*, 1994) using a polyclonal MrkA antibody. While no signal was detected in KP_ΔmrkA_, higher signal was seen in KP_mrkJ*_ compared to KP_WT_ (Fig. 2A). Flow cytometry analysis of live, fully capsulated KP cells revealed higher MrkA expression was observed in KP_mrkJ*_ with no detectable expression in KP_ΔmrkA_ (Fig. 2B). We also wanted to understand the impact of the mrkJ mutation on T3F configuration and subjected our strains to Western blot analysis. Once again, this confirmed higher MrkA expression in KP_mrkJ*_ but also indicated the presence of higher molecular weight (100 - 250 kDa) bands that were not denatured during SDS-PAGE (Fig. 2C). We excluded any impact any non-polar effects of MrkA expression levels on capsule and hypermucoviscocity by sedimentation assay (Fig. S2B). To confirm that enhanced MrkA expression in KP_mrkJ*_ is due to increased transcription we performed RT-qPCR. This revealed that *mrkA* expression was significantly higher in KP_mrkJ*_ compared to KP_WT_; expression of *mrkA* was not detected in KP_ΔmrkA_ (Fig. 2D).

**Figure 2.**
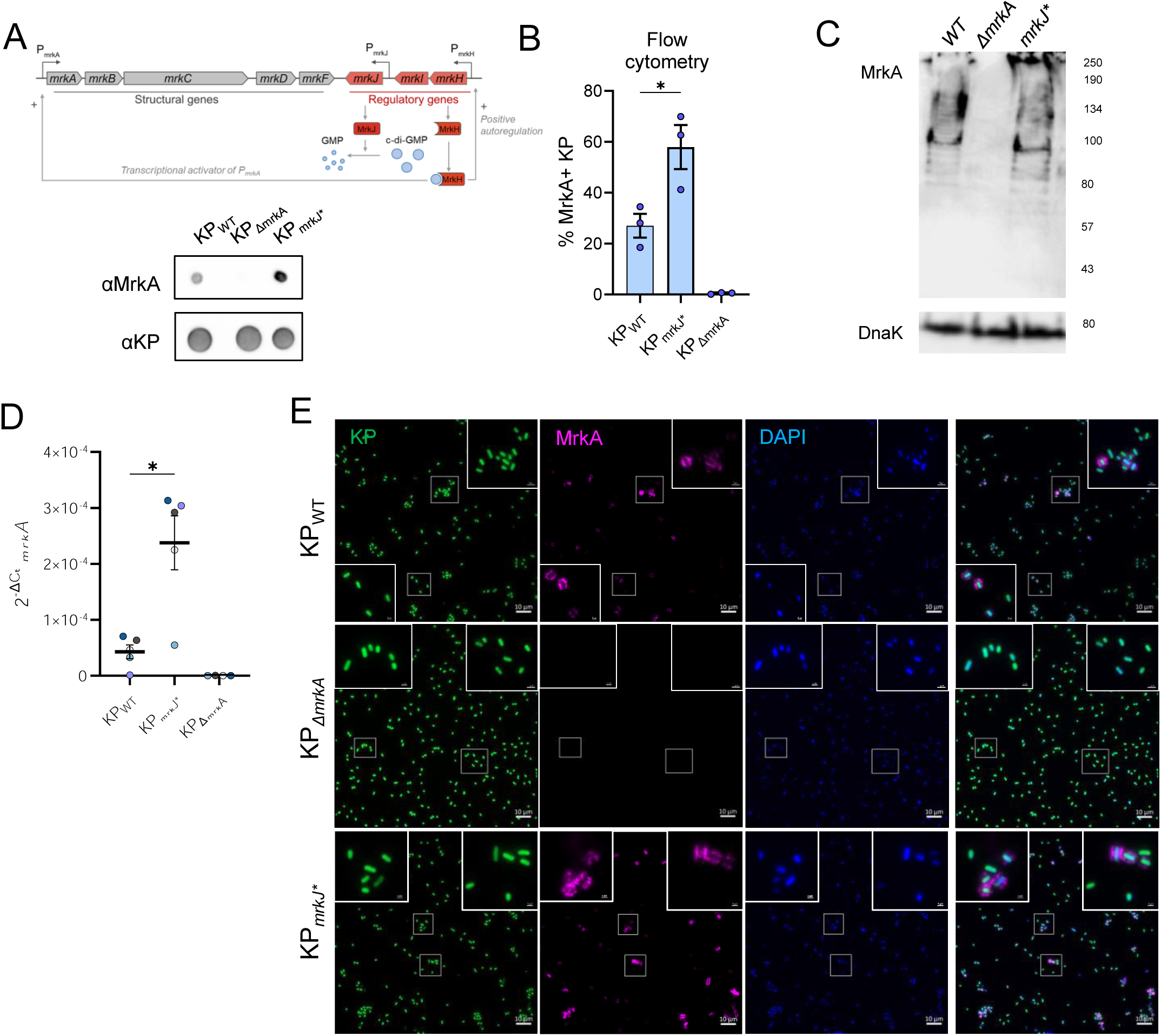
Development and Application of Phenotypic Assays to Study MrkA Expression Variability in KP A. (above) Simplified schematic illustrating the regulation of the *mrkABCDF* operon by *mrkHIJ*, where MrkH is an c-di-GMP-dependent activator and MrkJ functions as a repressor; the LuxR-type regulator MrkI also plays a positive regulatory role. (below) OD_600_ normalised whole cell samples for the indicated KP strains were spotted and probed with anti-MrkA antibody and anti-KP antibody was used as a loading control. The dot blot is representative of n=2 biological repeats. B. KP mutants grown in M9-Glc were stained with anti-MrkA primary antibody followed by fluorophore-conjugated secondary antibodies. The proportion of MrkA^+^ cells was measured for each strain using flow cytometry. The graph shows mean±SEM values for n=3 biological replicates; significance was determined using unpaired t test between KP_WT_ and KP_mrkJ*_. *, *P<0.05*. C. Whole-cell lysates from the indicated KP mutants expressing varying levels of MrkA were separated by SDS-PAGE and probed with anti-MrkA antibodies. Anti-DnaK was used as a loading control. Bands corresponding to polymerised MrkA (>80 kDa) were detected in KP_WT_ and KP_mrkJ*_, with expression levels increasing in KP_mrkJ_. Representative blot from n=3 independent experiments. D. RNA was extracted from indicated KP variants grown in M9-Glc, and expression of was quantified by RT-qPCR. Data were normalised to 16S rRNA as a housekeeping gene and analysed using the ΔCt method. Graph shows mean relative expression levels of mrkA from n=5 biological replicates, confirming differential transcription in the engineered mutants. Error bars represent standard error of the mean (SEM). Statistical significance was determined using Welch’s t test between KP_WT_ and KP_mrkJ*_. *, *P<0.05*. E. sfGFP-expressing KP strains were stained with anti-MrkA antibodies followed by Alexa Fluor 555–conjugated secondary antibodies. DAPI was used to stain DNA. Representative images show KP (green), MrkA signal (magenta) localised to the bacterial surface and DAPI (cyan). Scale bar = 10 µm. Images shown are representative of n=5-10 fields per mutant across n=5 biological replicates.

Finally, we used immunofluorescence (IF) staining of bacterial suspensions to visualise the T3F. This revealed specific T3F detection, which can be observed as punctate staining surrounding the T3F-expressing cells. Notably, KP expressing the T3F were frequently observed forming aggregates, which were not seen in KP_ΔmrkA_ (Fig. 2E). Taken together, these results confirm specific MrkA detection and that T3F expression is supraphysiological in the MrkJ mutant. These confirmatory data and techniques form the basis of the assays described below.

### Genotype-phenotype characterisation of MrkA in 100 strains *in vitro*

To investigate the effect of polymorphisms in the *mrk* operons on MrkA expression and the impact of a range of capsule types and mucoviscosity levels on MrkA binding by antibodies, we analysed a diverse curated collection of 100 KP strains (MRSN) (Martin *et al*, 2023). Using the short-read assembled genomes, we found that both the *mrkABCDF* and *mrkHIJ* operons were absent in MRSN21304, while MRSN560539 encodes a truncated copy of *mrkA* (Fig. 3A, denoted in orange with an asterisk). Moreover, analysis of the coding sequences showed that MRSN14444 (ST395) and MRSN740795 (ST4270) express MrkH variants G143S and R123L, respectively. These mutations affect the c-di-GMP and DNA-binding sites—both essential for MrkH’s function as a transcriptional activator (Johnson *et al*., 2011) (Fig. 3A, arrowheads in green, Table S3). We also identified fragmented copies of *mrk* genes in 4 additional strains: MRSN750877 (*mrk*C), MRSN562722 (*mrkD*) and MRSN375436 and MRSN730567 (*mrkH*) (Fig. 3A, Table S3). Among the 100 strains we observed 5 different MrkA protein level variants, 4 of which were identified as alleles 1, 2, 3 and 4 in our analysis of the EuSCAPE collection (Fig. 1C, Table S2), with the 5^th^ combining 2 SNPs present in EuSCAPE alleles 5 and 7. In line with our large genome collection analysis, MrkA alleles 1 and 2 accounted for 92% of the strains (Table S2 and S3). This also held true for MrkH where we observed 6 allelic variants, with alleles 1 and 2 from the EuSCAPE collection both present as the most prevalent MrkH sequences, accounting for 95% and 3% of the MRSN collection respectively (Fig. 1E, Table S2 and S3).

**Figure 3.**
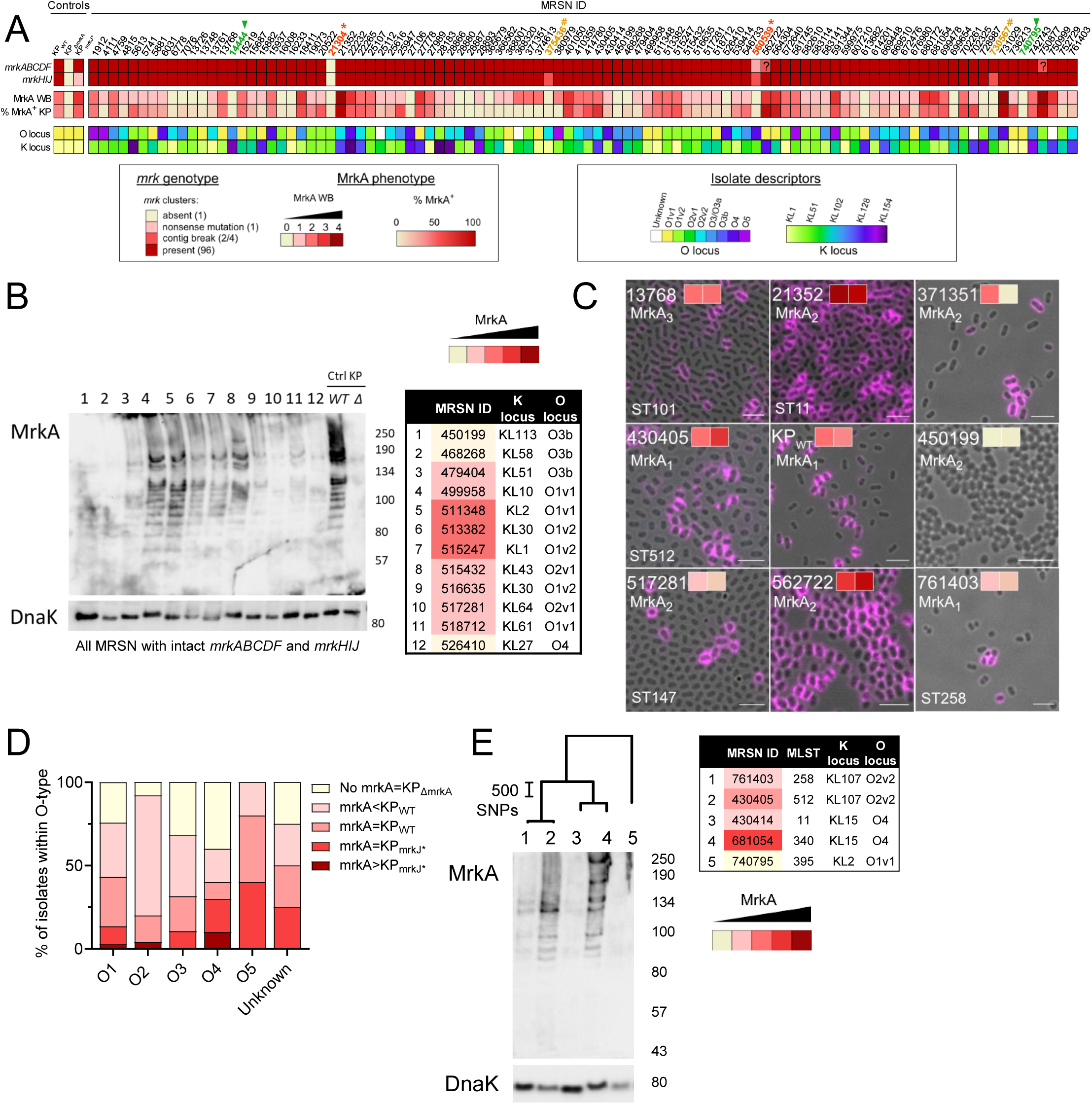
Linking Genetic Variation in *mrk* genes to Phenotypic MrkA expression using a collection of 100 representative KP Strains. A. Summary heatmap displaying the indicated features for the 100 *K. pneumoniae* isolates; KP_WT_, KP_ΔmrkA_ and KP_mrkJ*_ were used as controls; key is provided below. Columns represent individual isolates; rows represent features. Features displayed (top to bottom) are *mrkABCDF* prevalence and *mrkHIJ* prevalence (genotype), values obtained for mrkA expression by western blot and values obtained by staining in suspension and flow cytometry (phenotype). Additional data from Martin et al., 2023 is provided to reflect the variability: capsule (K) and O-antigen (O) loci assignments. 6 isolates with non-intact/ non-active mrkA and/or mrkH are highlighted: absent or truncated *mrkA* in orange with asterisk (*), key sites mutated in *mrkH* in green with arrowheads (▴) and fragmented *mrkH* in yellow with hashtag (#). Breaks observed in *mrkC* and *mrkD* are likely due to reading errors (denoted byin the heatmap box for the *mrkABCDF* cluster). Numerical values for the heatmaps and additional information can be found in Table S3, a representation of this data can also be found here: https://microreact.org/project/mrk-mrsn. B. Western blot analysis of polymerised MrkA protein expression across representative isolates indicated in the table; different shades of colour indicate different levels of MrkA detection. Bands were detected using anti-MrkA antibodies; lower panel shows the loading control DnaK. Blot is representative of n=3 biological repeats. C. Immunofluorescence microscopy images showing MrkA expression (magenta) in selected KP strains displaying various levels pf MrkA expression. MrkA expression phenotype (as determined by western blot and flow cytometry, from Fig. 3A) and MrkA allele are indicated for each panel. Scale bar = 2 μm. D. Relative abundance of MrkA expression phenotypes in strains belonging the indicated O-antigen types, binned into very high (higher than the MrkA-overexpressing control KP_mrkJ*_), high (equivalent to KP_mrkJ*_), medium (equivalent to KP_WT_), low (below KP_WT_) and undetectable groups across strains. Data is from the 94 stains from the collection with intact *mrkABCDF* and *mrkHIJ* genes. E. Western blot analysis for polymerised MrkA expression from selected MRSN isolates from the clonal complex (CC) 258 grown in M9-glucose. The branches showing how these strains are related is shown above the blot (scale=number of SNPs) and the table indicated the MRSN ID, ST and K (capsule) and O (O-antigen) loci, the colour indicates the level of MrkA expression; scale in phylogenetic tree represents the raw number of SNPs. The blot is representative of n=3 biological repeats.

We grew the 100 strains in the collection in M9-Glc and MrkA expression was characterised by Western blotting and flow cytometry; KP_WT_, KP_ΔmrkA_ and in KP_mrkJ*_ were used as controls (Fig. 3A and B, Fig. S3A, Table S3). MrkA was undetectable in the 4 strains with the *mrkA* and *mrkH* mutations identified above (MRSN21304, MRSN560539, MRSN14444, MRSN740795), validating the analysis, and in the 2 strains with fragmented *mrkH* genes (MRSN375436 and MRSN730567) (Fig. S3A, highlighted in different colours and symbols). Conversely, the two strains with fragmented copies of *mrkC* or *mrkD* both express MrkA, suggesting that the gene fragmentation is a result of a genome assembly failure rather than true disruption of the genes (Fig. 3A). The presence of the most prevalent MrkA and MrkH alleles in the EuSCAPE collection allowed us to confirm that the allele-specific SNPs, which have spread in circulating strains, do not directly affect the isolate’s ability to express MrkA (Figure 3A, 3C, Table S3). The levels of MrkA expression quantified by Western blot and flow cytometry correlated, independently of the levels of mucoviscosity, highlighting that antibodies can bind MrkA expressed by live KP even in the presence of a thick, mucoviscous capsule (Fig. S3B, Table S3). The level of MrkA expression in the 94 strains with intact *mrkABCDFHIJ* varied, independently of their *mrk* gene sequences, O and K types and their ability to form biofilm (Fig. 3A, 3C-D, Table S3). Furthermore, even phylogenetically related isolates, express different levels of MrkA (Fig. 3E, https://microreact.org/project/mrk-mrsn).

In order to further address whether these phenotypic differences can be predicted from the genomes, we also examined the upstream regions of *mrkABCDF* and *mrkHI* among the 94 isolates with intact *mrkABCDFHIJ*. Notably, no mutations affecting the RBS-10 and-35 boxes within the promoter sequences were observed (Tan *et al*., 2015; Wilksch *et al*., 2011) and only 4 SNPs <250bp away from the *mrkA* start codon were identified (SNPs described in Table S3). The upstream region of *mrkHI* was again more highly conserved, with 75% of isolates sharing an identical sequence, and the remaining isolates differing by only 1 or 2 SNPs (Table S3). Importantly, we found no association between upstream variation and the levels of MrkA expressed by MRSN isolates. Therefore, despite in-depth analysis of both operons and promoters, MrkA expression levels cannot yet be determined from genotype alone.

### The role of T3F in Pulmonary and Systemic Spread of KP

We next wanted to evaluate the impact of T3F expression on virulence. As pneumonia is a common primary focus for KP infections in patients we initiated infection in the lungs via intratracheal (IT) delivery with 500CFU of KP_WT_, KP_ΔmrkA_ or KP_mrkJ*_. We employed two mouse strains (BALB/c and CD-1) in which the kinetics of KP_WT_ infection are well defined, thus allowing us to introduce increased host diversity to more robustly assess the role of T3F in virulence. BALB/c mice deteriorate in a non-linear fashion after 24 h post infection (hpi) and reach endpoint at 36 hpi, whereas CD-1 infection exhibits delayed kinetics, with non-linear deterioration from 48 hpi with an end-point at 72 hpi (Wong *et al*., 2019; Wong *et al*, 2025). The latter better reflects the time course of early mortality in sepsis and septic shock (Daviaud *et al*, 2015).

In BALB/c mice all KP variants induced similar weight loss and increases in wet lung mass index, indicative of inflammatory lung infiltration and oedema, as well as lung CFU at 36hpi (Fig. 4A&B, S4A). Bacteraemia was present in 75%, 74.2% and 66.7 % of KP_WT_, KP_ΔmrkA_ or KP_mrkJ_ infected mice, albeit mostly at <10^5^ CFU/ mL, with no detectable differences between strains (Fig. 4C). We then proceeded to infect CD-1 mice and, at 72 hpi, KP_WT_ burdens in the lung averaged ∼2.5×10^9^, with all KP-infected CD-1 mice losing weight compared to mock-infected animals and displaying clear signs of lung damage and development of lobar pneumonia in highly infected mice (Fig. D&E, Fig. S4B). Blood bacterial burdens were higher in CD-1 infected mice compared to BALB/c mice, with a geometric mean more than 2-fold higher in the KP_WT_ group (Fig. 4F). Notably, while all KP_ΔmrkA_-infected CD-1 mice became bacteraemic, only ∼55% of the KP_mrkJ*_-infected mice had detectable blood CFUs at 72 hpi, with significantly higher systemic spread in KP_ΔmrkA_ compared to KP_mrkJ*_ infection (Fig. 4F). Correspondingly, there was a trend towards KP_ΔmrkA_ displaying higher levels of disease (Fig. 4D&E).

**Figure 4.**
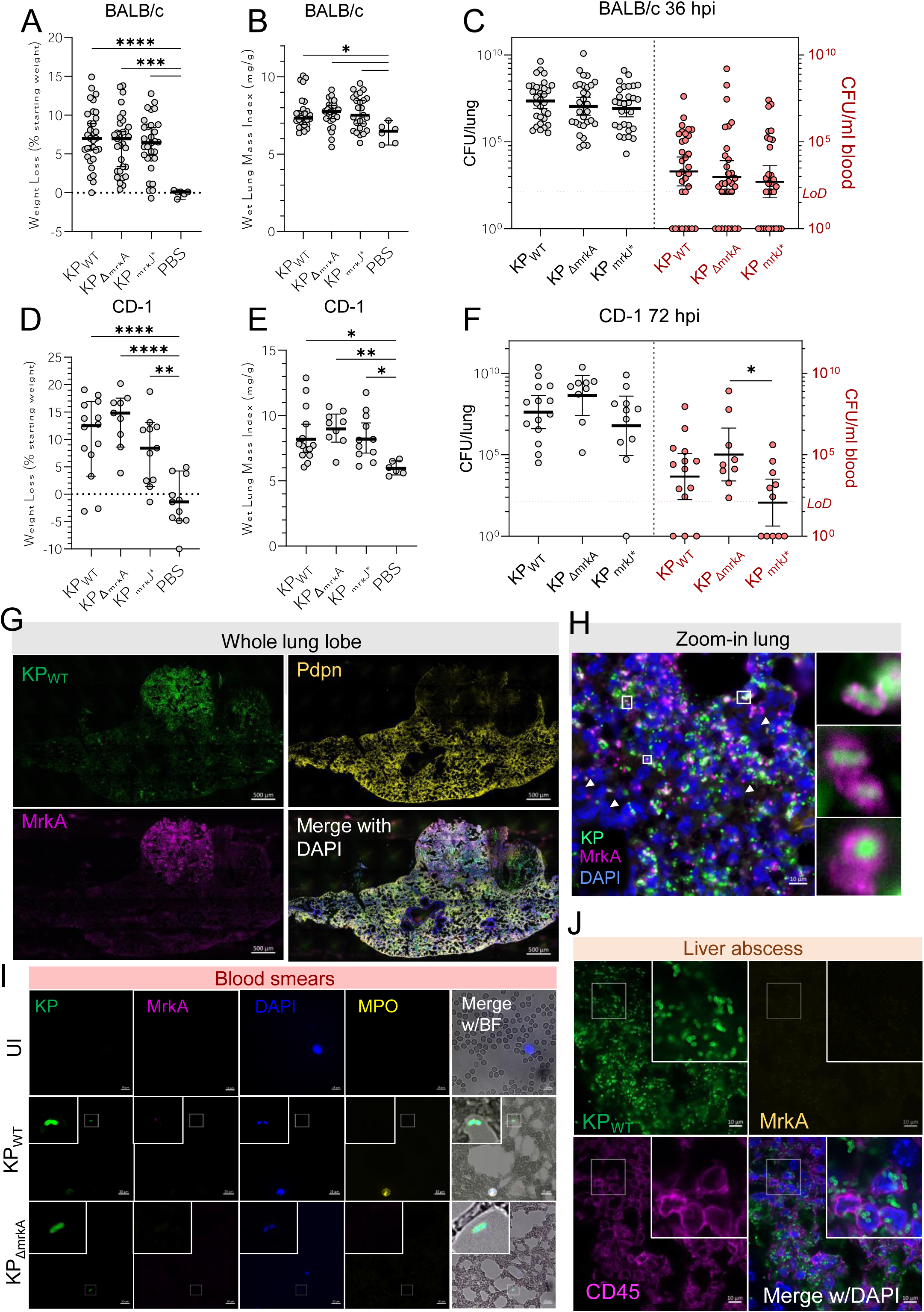
MrkA is detectable during KP lung infection and impacts systemic dissemination in a murine model of KP infection. A and B. BALB/c mice were infected by intratracheal infection with 500 CFU of KP_WT_, KP_ΔmrkA_ or KP_mrkJ*_; PBS was delivered as the uninfected control. At 36 hpi all infected mice lost weight (A) and displayed increased weight lung mass index (wet lung weight to body weight, B), associated to lung inflammation and oedema, which often correlates with histological evidence of consolidation and tissue damage. C. Bacterial burden in lungs and blood of KP-infected BALB/c mice at 36 hpi. Colony-forming units (CFU) per lung (left) and per ml of blood (right) are shown for each strain. Red dashed line indicates the limit of detection (LoD). D and E. CD-1 mice were infected by intratracheal infection with 500 CFU of KP_WT_, KP_ΔmrkA_ or KP_mrkJ*_; PBS was delivered as the uninfected control. At 72 hpi all infected mice lost weight (D) and displayed increased weight lung mass index (E) compared to the PBS control. F. Bacterial burden in lungs and blood of CD-1 mice at 72 hpi following infection with KP_WT_, KP_ΔmrkA_ and KP_mrkJ*_. Red dashed line indicates the limit of detection (LoD). G. Representative immunofluorescence staining of CD-1 lung tissue showing KP_WT_ (green) (1.48×10^9^ CFU/lung) and MrkA (magenta) co-staining in a consolidated area marked by the absence of podoplanin (Pdpn, yellow) as a marker for type I alveolar cells. DAPI is added as the nuclear counterstain in the merged imaged, highlighting the tissue context. Scale bar = 500 μm. H. High-magnification image showing KP_WT_ (green) and MrkA (magenta) colocalisation at the single-bacterium level in the lung; DAPI-stained nuclei display polymorphonuclear donut-like shape consistent with neutrophils (examples indicated by white arrowheads). Insets show enlarged views of individual bacteria expressing MrkA. Scale bar = 10 μm. I. Representative blood smear images from uninfected (UI) or infected (KP_WT_ or KP_ΔmrkA_) CD-1 showing no MrkA expression (magenta) in KP (green). DAPI (blue) stains host nuclei; MPO (yellow) highlights neutrophils which increase upon infection and brightfield overlay shows morphological context. Insets indicate zoomed views of bacteria. Scale bars = 10 μm. J. Immunofluorescence of liver tissue showing no MrkA expression (yellow), in an abscess with high levels of KP (green) and immune cells stained with CD45 (magenta); DPAPI stains host nuclei. Inset regions highlight KP clusters in proximity to or within immune cell–rich areas. Scale bar = 20 μm. (A-C, D-F) Graphs show mean±SEM (A, B, D, E) or geometric mean ± 95% CI from 5 (A-C) or 4 (D-F) biological repeats, each dot represents a mouse. Statistical significance was determined by ordinary one-way ANOVA (A, B, D, E) and lognormal ordinary one-way ANOVA (C, lung CFU and F, blood CFU), both corrected with Tukey’s multiple comparisons test and Kruskal-Wallis test with Dunn’s multiple comparisons test (C, blood CFU and F lung CFU) *, *P<0.05*; **, *P<0.01*; ***, *P<0.001*; ****, *P<0.0001*. (G-J) Images are representative of highly infected mice from n=4 biological repeats

The increased blood burdens of KP_ΔmrkA_ compared to KP_mrkJ*_ were not caused by increased sensitivity to complement-mediated killing or neutrophil phagocytosis (Fig. S4C). Similarly, phagocytosis assays with bone-marrow derived macrophages (BMDMs; as a proxy for monocyte-derived macrophages) showed that T3F did not affect KP uptake (Fig. S4D). When comparing bacterial burdens in the lung and blood, a clear correlation was observed in KP_ΔmrkA_-infected mice, with the lung CFU explaining 56% of the variance observed in blood CFU (*r^2^*=0.56). In contrast, this correlation was absent in KP_mrkJ*_-infected mice (*r^2^*<0.4), consistent with supra-physiological expression of MrkA impairing dissemination from the lung in this model (Fig. S4E).

### T3F Expression Varies Across Anatomical Sites

We wanted to understand MrkA expression levels during infection, including the primary site – the lungs – and secondary sites including the blood, liver and spleen. Immunofluorescence staining of lung sections revealed higher MrkA expression in areas of dense lung consolidation with decreased staining of the type I alveolar marker podoplanin, due to the local tissue destruction and influx of neutrophils denoted by their polymorphonuclear donut-like nuclei (Fig. 4G-H, Fig. S5A). Bacteria present in bronchiole lumen, delimited by E-cadherin-positive cells, displayed lower levels of MrkA expression (Fig. S5B). The MrkA staining pattern, when present, resembled hair-like protrusions encircling the bacteria (Fig. 4H). We noted that the MrkA expression appeared bimodal, consistent with the binary expression observed by flow cytometry (Fig. S3A) with KP existing as either MrkA positive (MrkA^+^) or negative cells (MrkA^-^) (Fig. 4H). We also prepared and stained blood smears from infected mice. Whilst KP_WT_ was readily detectable, no MrkA staining was observed (Fig.4I). Despite bacteraemia, we could not detect KP_WT_ in the liver of highly infected BALB/c mice at 36 hpi, while ∼10% of KP_WT_-infected CD-1 mice displayed formation of liver abscesses with detectable KP_WT_ (Fig. S5C-E). However, as observed in the blood no MrkA^+^ KP_WT_ were encountered in liver abscesses or in spleen, where KP_WT_ was found scattered in small clusters of up to ∼10 bacterial cells (Fig.4J, Fig. S5E-F). In summary we observed a non-uniform pattern of MrkA expression in the lungs during KP_WT_ infection and absent MrkA expression in the blood and metastatic foci of infection in distant organs. We then proceeded to quantitatively assess expression.

### Quantitative MrkA expression in the lungs and blood of infected mice

We developed a flow cytometry protocol to profile MrkA expression in vivo. Whilst MrkA was the target of these studies we aimed to develop a protocol that can be used to profile any antigen during natural *in vivo* infection, as long as specific detection reagent such as an antibody is available. To that end we infected mice with genomically-tagged sfGFP KP_WT_ and KP_mrkJ*_ and collected lungs and blood; mice inoculated with PBS or KP_ΔmrkA_ were used as uninfected and no MrkA controls respectively (gating strategy in Fig. S6A). To ensure that antigen levels were unchanged from their native state we fixed samples immediately following collection before downstream processing. We initially found that reliable quantification of MrkA expression in blood requires a minimum bacterial load of >9×10³ CFU/ml, which precluded robust analyses of MrkA expression in blood samples from BALB/c mice (Fig. S6B). However, CD-1 mice frequently develop bacterial loads in the blood exceeding 9×10³ CFU/ml (∼60% of KP_WT_ infected mice) (Fig. 4D) and proceeded with infections in this strain background for absolute quantification.

In the lungs, the proportion of MrkA^+^ KP_WT_ was ∼18%, while the proportion of MrkA^+^ in KP_mrkJ*_-infected mice was ∼31.5% (Fig. 5A-B, Fig. S6C). This confirmed that the nonsense mutation in *mrkJ* leads to a higher proportion of KP-expressing MrkA in vivo. In contrast, the proportion of MrkA^+^ KP_WT_ and KP_mrkJ*_ in the blood was ∼5% and ∼10% respectively (Fig. 5A, 5C). Performing paired comparisons with samples from the same mouse revealed a significant drop in the proportion of MrkA^+^ KP in the blood (Fig. S6D). This was consistent with our qualitative IF imaging and confirms that MrkA levels are lower in bacteraemic KP when the primary site of infection is the lungs (Fig 4I&J, S4&F).

**Figure 5.**
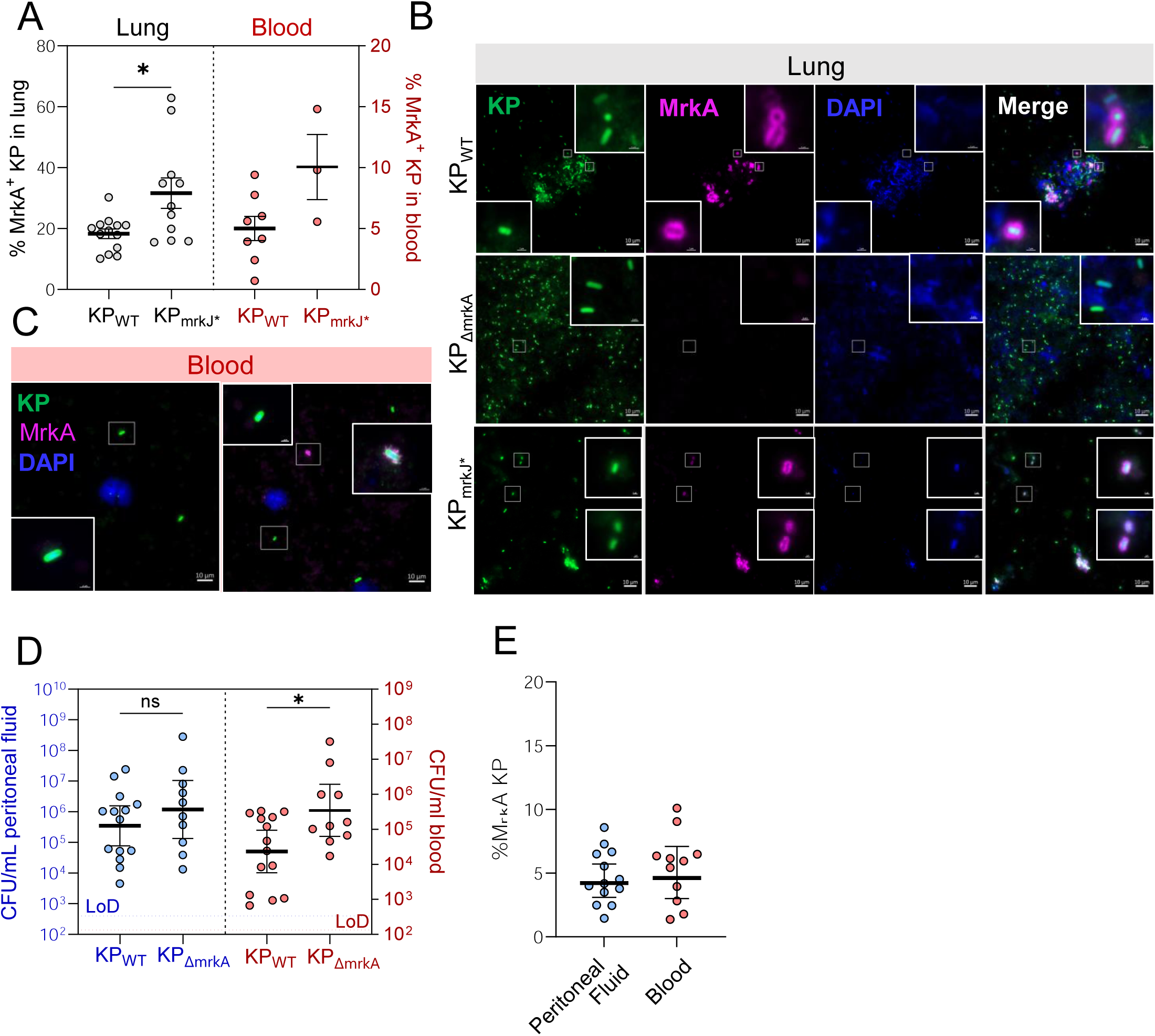
KP exhibits site-specific T3F expression patterns. A and B. Quantification of MrkA expression on KP (sfGFP⁺) isolated from the lung and blood, with higher levels shown by KP_mrkJ*_. Bacteria were recovered from infected tissue, stained for surface MrkA, and analysed by flow cytometry; uninfected and KP_ΔmrkA_ were used as controls to determine the KP and MrkA+ populations when gating (see Fig. S5A for gating). Lower levels of MrkA expression are observed in blood compared to lung and only mice with >4000 CFU/ml of blood were used for MrkA quantification. B. Lung samples from mice infected with the indicated KP strains expressing sfGFP (green),stained with anti-MrkA (magenta) and DAPI (dark blue) and imaged using agarose pads. KP_WT_ and KP_mrkJ*_ show positive surface staining of MrkA and distinct clumping of the bacterial cells while no MrkA is observed in KP_ΔmrkA_, which are more evenly distributed. Images are representative of 4 biological repeats. Scale bar = 10 µm. C. Blood samples from KP_WT_ infected mice; the image on the left shows KP (green) with no MrkA, while the one of the right has one bacterium expressing MrkA (magenta), shown in the inset, which was rare to find. DAPI was used to stain the nuclei. Images are representative of 4 biological repeats. Scale bar = 10 µm. D. Bacterial burdens in peritoneal fluid and blood from CD-1 mice infected with the indicated strains via the intraperitoneal route. Samples were collected at 48 hpi; KP_ΔmrkA_ bacteraemia is significantly increased in the blood after intraperitoneal inoculation compared to KP_WT_. E. Quantification of MrkA expression on sfGFP-expressing KP_WT_ isolated from the peritoneal fluid and blood reveals similar low levels of expression. Bacteria were recovered from infected tissue, stained for surface MrkA, and analysed by flow cytometry. A, D, E. Each dot represents measurements obtained from an individual mouse; graphs show mean±SEM (A, E) or geometric mean± 95% CI from n=4 (A), n=3 (E) or n=3 (KP_WT_) or 2 (KP_ΔmrkA_) (D) biological repeats. s with statistical significance determined by unpaired Welch’s t test (A), lognormal or unpaired t test (D). *, *P<0.05*.

To further evaluate if T3F expression levels impact on systemic spread and if low MrkA KP expression is a feature of KP in circulating blood we used an alternative model-intraperitoneal (IP) infection. Here KP initiates infection in peritoneal cavity before disseminating into the bloodstream and we infected CD1 mice with KP_WT_ and KP_ΔmrkA_. Consistent with lung infection, no differences in peritoneal lavage fluid (PL) CFUs were observed, however, higher dissemination to the blood occurred in KP_ΔmrkA_ compared to KP_WT_ (Fig. 5D). The proportion of MrkA^+^ KP_WT_ in the blood and PL was ∼5%, similar to the proportion of blood MrkA^+^ KP_WT_ following dissemination from lung infection (Fig. 5E). Taken together, these results indicate that a minor population of KP express MrkA in the lungs, which further drops when KP enters the blood and that MrkA expression by KP in the blood is low, irrespective of primary focus. Interestingly our data also suggest that MrkA expression in the lungs itself may limit systemic KP dissemination.

### Antigen Abundance Modulates mAb Efficacy in KP Infection

As MrkA has been proposed as a target for mAb treatment, we evaluated the efficacy of KP3 administration (Wang *et al*., 2016) (a MrkA binding human mAb) in the context of physiological (KP_WT_) and supraphysiological T3F expression (KP_mrkJ*_). To test this, we used the BALB/c infection model (Fig. 4B) as we found indistinguishable levels of bacterial replication in the lungs and dissemination to the blood between the two mutants in this shorter time course. KP3 mAb, or an isotype human IgG mAb control, were administered 24 h before infection with KP_WT_ or KP_mrkJ*_ (Fig. S6E). While KP3 administration did not affect infection induced weight loss, it significantly reduced lung damage (wet lung mass index) specifically in KP_mrkJ*_-infected mice (Fig. 6A, S6F). Moreover, while KP3 resulted in reduced lung CFUs in both KP_WT_ and KP_mrkJ*_ infection it only significantly reduced bacteraemia in KP_mrkJ*_ infection (Fig. 6B-C). Serum inflammatory markers, such as TNF, IL-6 and CXCL1, were also reduced, but only significantly in KP_mrkJ*_-infected mice (Fig. 6D-F) Finally, we quantified serum creatinine, a marker of acute kidney injury, one of the most common complications of sepsis (Akcan Arikan *et al*, 2025). This revealed that while KP3 reduced serum creatinine in KP_WT_-infected mice, the effect only became significant in the KP_mrkJ*_-infected mice (Fig. 6G). These results suggest that increased MrkA expression boosts the efficacy of KP3 and significantly reduce infection related sequalae. Therefore, as the level of antigen expression correlates with mAb prophylactic efficacy, this parameter should be considered in developing immunotherapeutics targeting bacterial infections.

**Figure 6.**
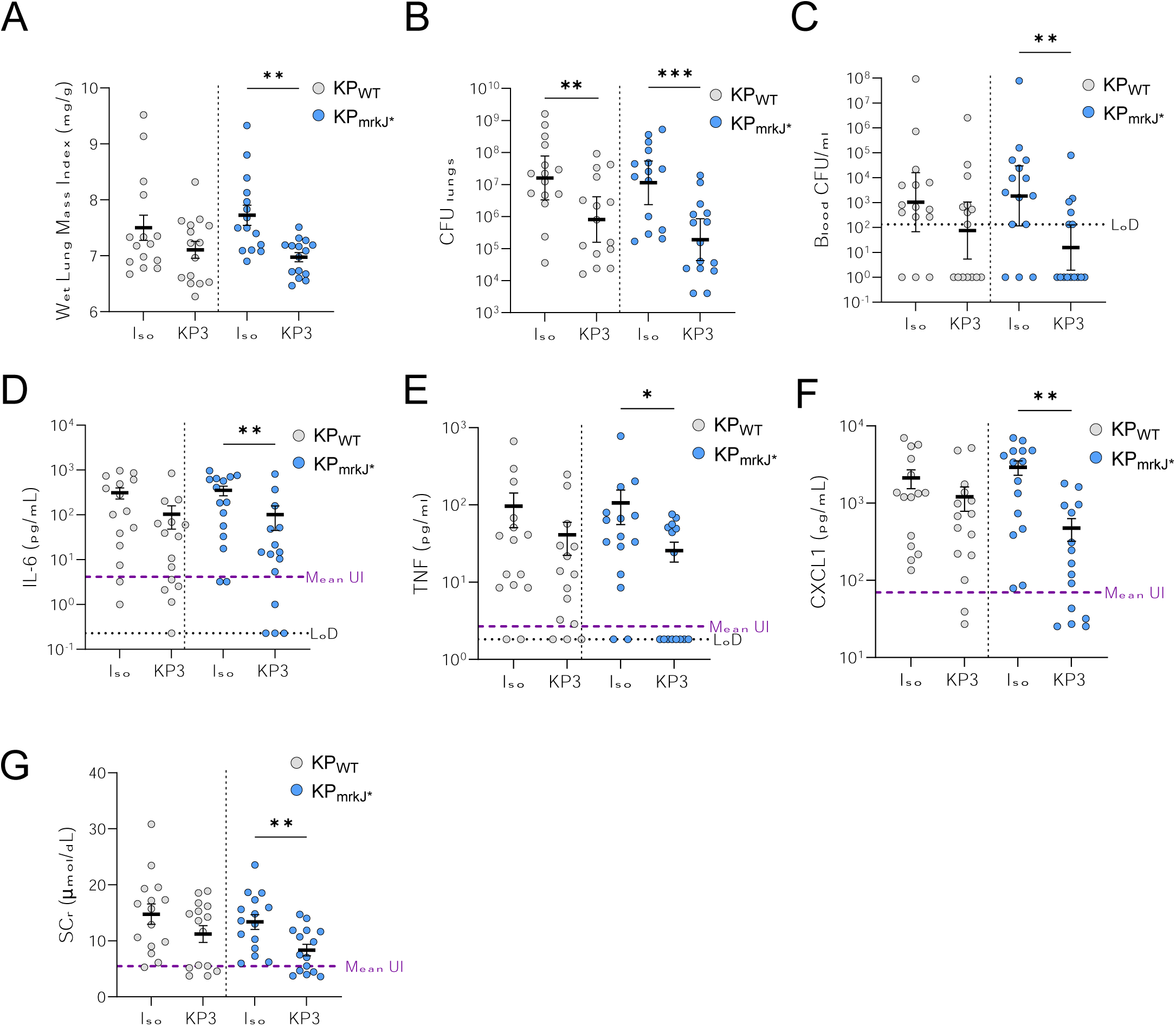
T3F expression directly influences the effectiveness of monoclonal antibody-based interventions. BALB/c mice were prophylactically administered anti-MrkA mAb KP3 or isotype control (15 mg/kg) and challenged by intratracheal infection with 500 CFU of KP_WT_ or KP_mrkJ*_ 24 h after. At 36 hpi we analysed the following indicators of infection and disease severity, which indicated increased effectiveness of KP3 administration in treating infection with the KP variant that overexpresses MrkA, KP_mrkJ*_. A. KP3 prophylaxis in KP_mrkJ*_-infected mice significantly prevents the increase in wet lung mass index (wet lung weight to body weight) associated to lung inflammation and oedema. B and C, Bacterial replication in lung (B) and blood (C) were reduced in mice administered KP3 but the effect was more significant in KP_mrkJ*_-infected mice. D-F. Serum levels of the pro-inflammatory cytokine IL-6 (D), TNF (E) and the neutrophil chemoattractant CXCL1 (F) are improved by KP3 administration compared to the isotype control. Similar to what is seem with the wet lung mass index, a significant reduction is only seen in KP_mrkJ*_ infected mice. This indicates that increased expression of the antigen increases the efficacy of the mAb. A black dotted line shows the limit of detection (LoD) for TNF and a purple dashed line indicated the mean value for uninfected (UI) mice. F. Acute kidney injury and impaired renal function are reduced in KP3-treated mice compared to those administered isotype control, but the trend is significant only in KP_mrkJ*_ infection. Graphs show mean±SEM (A, D, E) or geometric mean ± 95% CI (B, C) from n= 3 biological repeats. Each dot represents an individual mouse; the purple dotted line represents the mean value obtained from mock-infected mice. Statistical significance was determined by unpaired Welch’s t test (KP_mrkJ*_ in A); t test (KP_WT_ in E and F, KP_mrkJ*_ in G), Mann-Whitney test (KP_WT_ in A and G, KP_mrkJ*_ in E and F); Lognormal t test (KP_WT_ in B and C, both in D) and Lognormal Welch’s t test (KP_mrkJ*_ in B and C). *, *P<0.01*; ***, *P<0.001*.

## Discussion

Bacterial infections are becoming difficult to treat due to increasing rates of antibiotic resistance, leading to high mortality and significant healthcare burdens, with KP becoming one of the top-priority pathogens for global public health agencies (Sati *et al*, 2025). For alternative treatments such as vaccines (WHO, 2024b) and mAbs to be effective, it is paramount to choose an antigenic target that is not only prevalent and conserved in circulating pathogens, but also expressed during infection. While limited studies have addressed this question in vivo, it has been previously shown that different disease contexts can impact the effectiveness of mAb targets, such as the KB001 mAb candidate against the *Pseudomonas aeruginosa* T3SS which showed efficacy in ventilator associated pneumonia but not in cystic-fibrosis patients (François *et al*, 2012; Jain *et al*, 2018).

One of the most highly studied KP antigen candidates is MrkA, the major component of T3F (Berry *et al*., 2024; Wang *et al*., 2016; Wang *et al*., 2017). The main objective of this study was to determine its potential as a therapeutic target by performing an in-depth characterisation of its prevalence, conservation, epitope accessibility and levels of expression both in vitro and during in vivo infection. The workflow described here for KP and MrkA could be applied to future pathogen/antigen combinations with the only constraint being the availability of a specific antigen detection reagent (Synopsis Figure).

Analysis of large clinical genome collections revealed that the *mrkABCDF* operon is prevalent, agreeing with previous smaller studies (Bialek-Davenet *et al*., 2014; Gual-de-Torrella *et al*, 2022; Onori *et al*., 2015; Vuotto *et al*., 2017), and its genes are predominantly intact. The *mrkHIJ* regulatory operon is also conserved in KP, including widely used laboratory strains, with Kp52145 and some ST258 isolates being the exception due to IS-mediated disruption of *mrkH* (Becker *et al*, 2018).

We harnessed the diversity provided by the MRSN collection to enable us to investigate MrkA expression and confirmed that MrkA can be bound by antibodies in the presence of thick capsule, a notable finding given that this structure has been found to limit the accessibility of other surface antigens (Wantuch *et al*, 2024). T3F are thought to be the main contributing factor to biofilm formation in KP and overall, all biofilm producing strains were able to express MrkA. However, four MRSN isolates, 16008, 27106, 28880 and 450199, are capable of forming biofilms (Beckman *et al*, 2024) without MrkA expression, pointing towards alternative mechanisms such as type 1 fimbrial (*fim)*-dependent biofilm formation (all isolates are from urine samples and encode *fimH*) (Beckman *et al*., 2024; Rosen *et al*, 2008), PNAG production (both 27106 and 28880 express PNAG) (Bradshaw *et al*, 2024) or differences in the cellulose *bcs* operon (Zogaj *et al*, 2001).

These results highlight the need to look beyond the simple presence of intact structural genes and extend surveillance to the regulatory gene *mrkH*. Indeed, we show that an insertion sequence in *mrkH* abolishes *mrkA* expression, even in the presence of the other components of the operon. Moreover, study of upstream sequences revealed high conservation of the promoter sequences, except for an insertion sequence that appears to disrupt the *mrkH* promoter in some ST101 isolates. These strains have clonally expanded in a significant proportion of this high risk ST, a cKP with increasing presence which may not require T3F. This implies that conservation of upstream sequences should also be established during studies of promising antigenic targets.

Nonetheless, profiling of MrkA expression in our isogenic mutants shows that, although the overall proportion of bacteria expressing T3F is increased in the KP ^⁎^ mutant, expression within the population remains bimodally distributed with the highest proportion remaining MrkA^-^. This is consistent with the observations of Berry *et al*. and shows that T3F expression cannot currently be determined from genomic sequence of the structural *mrkABCDF* operon alone (Berry *et al*., 2024). Furthermore, we bolster these findings by profiling MrkA expression in the MRSN collection where we find a poor correlation between genomic sequence and phenotypic expression.

MrkA profiling revealed that ∼20% of KP in the lung express MrkA, with higher expression noted in consolidated alveolar spaces exhibiting large accumulations of neutrophils. This indicates that KP can form aggregates in the lungs which could impact antibiotic effectiveness, and could be reduced by anti-MrkA administration (Wang *et al*., 2016). However, the proportion of MrkA^+^ bacteria drops in the blood to ∼5%, whether this active bacterial adaptation of this environment remains to be shown. Staining of liver abscesses and spleen showed similar results, agreeing with previous studies where KP in systemic sites were shown to be more similar to each other than the lung (Holmes *et al*, 2025). These results, along with the conservation of both structural and regulatory sequences observed in clinical isolates, suggest that tight regulation of T3F expression is a critical feature of KP infection. It is likely that T3F are initially upregulated to support proliferation within a specific niche, and subsequently downregulated as nutrient availability declines. For instance, glucose (Lin *et al*, 2016) and iron (Chu *et al*., 2023; Wu *et al*, 2012) have been shown to positively regulate T3F expression.

The effectiveness of our KP3 mAb therapy reflected our MrkA profiling, clearly establishing that higher antigen expression levels result in better mAb activity. Our in vivo results predict reduced effectiveness of anti-MrkA mAbs in primary KP blood stream infections. However, they could still be effective in the initial stages of pneumonia and the reduction in blood burdens we see in our mAb prophylaxis model are likely due to a reduction in lung CFUs. Further, previous work has shown T3F to be required in catheter-associated urinary tract infections (Murphy Caitlin *et al*, 2013) and for gut colonisation (Tu *et al*, 2009), where this antigen may be more highly expressed and mAb therapy and/or vaccines may prove more effective here.

In the future, comprehensive profiling of antigen prevalence, conservation and expression both in vitro and in vivo as we have done for MrkA (Synopsis Figure) will ultimately serve to inform the design of effective prophylactic and therapeutic strategies. This approach allows resource allocation into developing the treatments that have the most potential and implementing them in the populations that will benefit the most.

## Methods

### Genome collections

We investigated the presence and variation among the *mrkABCDF* and *mrkHIJ* genes and upstream regions using two collections of publicly-available genomes. The first comprised 1717 short-read assembles from a European-wide collection of clinical *K. pneumoniae* species complex isolates (David *et al*., 2019). The second comprised short-read assemblies obtained from a panel of 100 diverse *K. pneumoniae* from the Multidrug-Resistant Organism Repository and Surveillance network (MRSN) (https://pathogen.watch/collection/8zjjb34v3vh9-mrsn-diversity-panel) (Martin *et al*., 2023).

### Identification and characterisation of *mrkABCDF* and *mrkHIJ* among public genomes

To identify each gene from the genome assemblies, we used BLASTn v2.14.1 (Camacho et al. 2009) using a query gene sequence from the reference genome, ATCC43816 (accession CP009208). We required single hits per assembly that matched ≥25% of the query length and contained a start codon. The resulting nucleotide sequences were aligned using MUSCLE v5.1.0 (Edgar, 2004). To assess variation among the encoded proteins, the nucleotide sequences were also translated to amino acid sequences using Seaview v5.0.5 (Gouy *et al*, 2021) using the standard genetic code. Protein sequences were considered truncated if a stop codon was identified within the first 90% of the query sequence length. The gene and protein variants identified among the European-wide collection were analysed within a phylogenetic context using a core gene-based phylogeny generated previously (David *et al*., 2019) via Microreact (Argimón *et al*, 2016). We also identified the upstream regions of *mrkA* (376bp) and *mrkH* (698bp) among the genome assemblies using a similar approach, requiring hits that matched ≥75% of the query length (with a higher threshold used to avoid a large abundance of short matches).

### Bacterial cultures

Bacteria were routinely cultured in LB Broth (Miller) or M9 minimal media supplemented with 0.4% glucose (M9-Glc). Saturated (overnight, 18 hour) cultures took place by inoculating broth and incubation at 37°C and shaking at 200rpm.

Antibiotics were added to LB broth or LB agar at the concentrations detailed in Table S4.

### Generation of isogenic strains

The homologous recombination technique resulting in scarless markerless mutants was used as described previously (PNAS 2022) to generate all strains based on the Standard European Vector Architecture platform pSEVA612S (JX560380.2) plasmid (https://seva-plasmids.com/).

pSEVA612S vectors for mutagenesis were generated by routine molecular biological techniques, including genomic DNA extraction, polymerase chain reaction (PCR) and Gibson assembly, plasmid extraction and PCR purification, all conducted following the manufacturer’s instructions (all reagents and kits in Table S4). Custom primers in this study were made by Merck (Sigma-Aldrich) and detailed in Table S4. Sanger sequencing of plasmids, to check vector construction, and PCR products, to asses seamless integration into the genome, were carried out by Eurofins Genomics.

The genomic DNA reference sequence used for ICC8001 is CP009208.1, obtained from the closely related ATCC43186 KPPR1. A list of strains and plasmids used and generated in this work can be found in Table S4.

### Dot blot

Bacterial cultures were grown overnight in M9 medium supplemented with glucose (M9-Glc), normalised to an OD₆₀₀ of 1.0, and centrifuged to pellet the cells. Pellets were resuspended in 1 mL PBS, and equal volumes were spotted onto a nitrocellulose membrane (#88018, ThermoFisher Scientific), then air-dried. Membranes were blocked for 2 hours at room temperature in 5% milk in PBS-T, followed by overnight incubation at 4°C with anti-MrkA polyclonal antibody (pAb) diluted in 5% BSA in PBS-T. After three washes in PBS-T, an HRP-conjugated anti-rabbit secondary antibody (see Table S4) diluted in 5% milk in PBS-T was applied and incubated for 1 hour at room temperature, followed by three additional washes. Signal detection was performed using ECL substrate (Bio-Rad), and images were captured using the ChemiDoc XRS+ system and analysed with Image Lab software (Bio-Rad Laboratories).

### Western Blotting

Bacterial cultures were grown overnight in M9-Glc and centrifuged to pellet the cells. Samples were lysed in Qubit-compatible lysis buffer (2% SDS, 1% glycerol, 60mM Tris pH 6.8) for 10 mins, boiled and vortexed; protein concentrations were then determined using a Qubit 4 Fluorometer (ThermoFisher Scientific). Lysates were then mixed with 5x Laemmli loading buffer (50 mM Tris pH 6.8, 2 % (w/v) SDS, 10 % (v/v) glycerol, 0.01 % (w/v) bromophenol blue, 0.05 % (w/v) 2-mercaptoethanol). Samples were boiled and 15 µg of protein sample were separated by SDS–polyacrylamide gel electrophoresis (PAGE) using a 7% Acrylamide/Bis-acrylamide gel (37.5:1) and tris-glycine buffer system. Samples were transferred to 0.2 µm polyvinylidene difluoride membranes (#1620177, Bio-Rad Laboratories) using a TransBlot semi-dry electrophoretic transfer machine (Bio-Rad) and a discontinuous Tris-CAPS Buffer System where the stock buffer is composed of 60 mM Tris and 40 mM CAPS, pH 9.6, and is supplemented with either 15% methanol (anode) or 0.1% SDS (cathode). Membranes were blocked for 2 h at room temperature in 10% fat-free milk PBS–0.1% Tween 20 (PBS-T) and incubated overnight at 4°C with anti-MrkA pAb in PBS-T with 5% BSA. Secondary antibodies were added in 5% fat-free milk in PBST and incubated for 1 h at room temperature. Immunoblots were developed with Clarity Western enhanced chemiluminescence (ECL, Bio-Rad Laboratories) and images were acquired on a ChemiDoc XRS+ and visualised using Image Lab software.

### Bacterial cell immunofluorescence microscopy

The indicated bacterial strains were grown overnight in M9-Glc. A 500 µL aliquot of the overnight culture was used for staining. Cells were pelleted by centrifugation at 5000 × g for 5 minutes 4 °C and washed two times with PBS. After establishing that fixing with paraformaldehyde (PFA) did not alter the MrkA staining pattern, we routinely followed the washed with incubation in 1% PFA/PBS for 20 mins on ice to prevent any changes in mrkA expressing during the antibody incubation periods. Fixing was followed by two washed in PBS and blocking for 20 mins in 5% BSA/PBS. Primary staining with anti-MrkA antisera was performed overnight at 4 °C in 1% BSA/PBS with agitation. After staining, cells were washed two times with PBS using centrifugation (5000 × g, 5 minutes, 4 °C) and resuspension. Secondary antibody and, if indicated, DAPI were added in 1% BSA/PBS and incubated for 45 minutes at room temperature with agitation. Cells were again washed three times in PBS using the same centrifugation and resuspension steps. Finally, cells were washed once more and resuspended in 50 µL of fresh PBS for imaging. 5 µl of the final preparation were imaged by means of a freshly made 2% agarose in (Milli-Q water) pad, using a Zeiss AxioImager Z1 microscope with a Hamamatsu microscope camera and processed using ZEN3.1 software (ZEN lite, Carl Zeiss MicroImaging GmbH, Germany).

### MrkA quantification from cultures by flow cytometry

The KP MRSN Diversity Panel (NR-55604) was obtained from BEI Resources, NIAID, NIH (Martin 2023). Isogenic Klebsiella pneumoniae strains— KP_WT_, KP_ΔmrkA_ and KP_mrkJ*_ — expressing sfGFP were grown to saturation overnight in M9-glc (see Fig. 2). Alternatively, the 100-strain MRSN collection, along with control strains (KP_WT_, KP_ΔmrkA_ and KP_mrkJ*_), were cultured overnight in 0.8 mL M9-glc in deep-well 96-well plates (#732-3802, VWR) in a ping-pong incubator. Following overnight growth, 20% of each culture was transferred to a V-bottom plate (Greiner #651201) for staining. Cultures were centrifuged at 4000 rpm for 10 minutes at 4°C, washed once in PBS, and fixed in 1% paraformaldehyde (PFA) in PBS for 20 minutes at 4°C using a microplate mixer (IKA-3208001). Samples were then centrifuged and washed twice in PBS, resuspended in 5% BSA/PBS, and blocked for 1 hour at room temperature with shaking.

After blocking, samples were centrifuged and resuspended in anti-MrkA polyclonal antibody (pAb) diluted in 1% BSA/PBS to be stained overnight at 4°C with shaking. The next day, samples were washed twice in PBS and incubated with Alexa Fluor-conjugated anti-rabbit IgG (see Table S4) for 1 hour at room temperature, in the dark and with shaking. Finally, samples were washed twice more in PBS and stored in the dark until flow cytometry. Flow cytometric acquisition was performed on a Cytek Amnis CellStream, analysing a minimum of 50,000 bacterial cells per sample. Data analysis was carried out using FlowJo v10.10.0.

### RNA isolation and reverse transcription quantitative PCR (RT-qPCR)

The indicated KP strains were cultured overnight in M9-Glc at 37 °C shaking. A 1 mL aliquot of each overnight culture was washed once with PBS and incubated with RNAprotect Bacteria Reagent for 10 minutes at room temperature. Cells were then pelleted by centrifugation at 5000 × g for 5 minutes. The resulting bacterial pellet was resuspended in 100 µL of TE buffer (30 mM Tris-Cl, 1 mM EDTA, pH 8.0) containing 15 mg/mL lysozyme and 20 µL of Proteinase K, and incubated for 10 minutes to lyse bacterial cells. 350 µl RLT lysis buffer supplemented with 2-mercaptoethanol was added to each sample and thoroughly mixed to ensure complete lysis. RNA was then extracted using the RNeasy Mini Kit (Qiagen) in accordance with the manufacturer’s instructions. RNA concentration and purity were assessed using a Nanodrop 2000 spectrophotometer (ThermoFisher Scientific).

To remove genomic DNA, 1 µg of the extracted RNA was treated with 1 U of RQ1 RNase-free DNase at 37 °C for 1 hour followed by heating for 10 minutes at 65°C in the presence of Stop Solution to inactivate the enzyme. cDNA synthesis was carried out using 200 U/reaction of M-MLV Reverse Transcriptase with random primers, as per the manufacturer’s protocol. A no-reverse-transcriptase (N-RT) control was included to confirm the absence of residual genomic DNA.

Quantitative PCR (qPCR) was performed using PowerUp SYBR Green Master Mix and validated primers with efficiencies between 95–105% (see Table S4 for the sequences). Reactions were run on a QuantStudio 1 System (Applied Biosystems, ThermoFisher Scientific), and data were analysed using QuantStudio Design & Analysis Software. Relative mRNA expression levels in each sample were calculated using the 2–ΔCt method, with 16S rRNA serving as the internal reference gene.

### Neutrophil isolation from the bone marrow (BM)

CD-1 mice were euthanised by cervical dislocation and tibia, femur, and hip bones were extracted and cleaned to then be crushed in PBS with 2 % FBS/PBS. The cells were filtered through a 40-µm strainer and centrifuged 300 xg for 10 min at 4°C. Pellets are resuspended in red blood cell (RBC) lysis buffer (10 mM potassium bicarbonate, 155 mM ammonium chloride, 0.1 mM EDTA, and 5 % FBS in water) and incubated for 5 mins at room temperature. After a second centrifugation at 4 °C, cells were resuspended in MACS buffer (50 mg/ml BSA, 2 mM EDTA in PBS) and counted using a haemocytometer; viability was checked via trypan blue exclusion (∼95% viability). Neutrophils were then enriched using anti-Ly6G microbeads following the manufacturer’s protocol, using LS MACS columns (#130-042-401) and a quadroMACS separator (all from Miltenyi Biotec). Purity was determined by flow cytometry and was shown to be 96 - 99 % Ly6G^+^ neutrophils. Neutrophil infections were performed on the same day as the extraction as detailed below.

### Differentiation of primary BMDMs

Tibia, femur, and hip bones were extracted from euthanised CD-1 mice and cleaned in DMEM containing gentamicin to then be crushed in PBS with 2 % FBS/PBS. The cells were filtered through a 70-µm strainer, centrifuged 300 *xg* for 10 min at 4°C and seeded into non-tissue culture treated 10 cm petri-dishes in DMEM containing 4500 mg/l glucose, 10% heat-inactivated fetal bovine serum (HI-FBS) and 2 mM GlutaMAX (cDMEM), supplemented with 50% spent media from L929 cells (sL929; containing M-CSF) and 100 U/mL penicillin and 0.1mg/mL streptomycin (PS). On day 2 of differentiation into bone marrow-derived macrophages (BMDMs), media was replaced with fresh cDMEM with 50% sL929. On day 4 of differentiation into BMDMs, adherent cells were passaged if confluent, progressively reducing the concentration of sL929 to a final concentration of 20% between passages and media changes. Cell detachment was performed by incubation with cold DPBS-2mM EDTA. Primary BMDMs were maintained in cDMEM supplemented with 20% sL929 and were used for infection as detailed below between 7-15 days post-extraction.

### *In cellulo* infection and antibiotic protection assay

Neutrophils were seeded in RPMI in 96-well plates at 2 × 10⁵ cells per well and incubated at 37°C in a humidified incubator with 5% CO₂ for 1 h until they attached to the bottom of the plate before phagocytosis assays were carried out. BMDMs were seeded in cDMEM into 48-well plates at a density of 4 × 10⁵ cells per well the day prior to infection.

Indicated KP isogenic strains were grown overnight and added to cells at a multiplicity of infection (MOI) of 10, confirmed by retrospective plating. Bacterial density was estimated by measuring optical density at 600 nm (OD₆₀₀), where an OD₆₀₀ of 1 was considered equivalent to 9 × 10⁸ CFU/mL. To synchronize infection, plates were centrifuged at 1000 rpm for 5 minutes at room temperature, followed by incubation for 45 mins at 37°C in 5% CO₂. Extracellular bacteria were then killed with the addition of gentamicin (200 µg/ml final) for 15 mins. At the end of the infection, cells were washed three times with sterile PBS and eukaryotic sells were lysed in sterile 0.5% triton X-100/PBS, serially diluted in PBS and plated in LB agar containing rifampicin. Plates were incubated overnight at 37°C and colonies were counted the following day.

### Serum Bactericidal Assay

Overnight bacterial cultures grown in M9-Glc were subcultured 1:20 in fresh M9-Glc, and cultures were incubated at 37 °C, shaking for 1.5 h to reach early logarithmic growth phase. Bacterial density was estimated by measuring OD₆₀₀ as above, and cultures were centrifuged for 3 min, and the pellets were resuspended in sterile PBS to achieve a final concentration of approximately 2 × 10⁶ CFU/mL.

For the serum bactericidal assay, 80 μL of the bacterial suspension was mixed with 80 μL of either pooled normal human serum (NHS) or heat-inactivated serum (56 °C, 30 min) in 96-well U-bottom plates (Falcon, SLS), in duplicate. An initial 30 μL aliquot was collected from each well immediately after mixing (0-h timepoint), serially diluted, and plated on LB agar plates. The microtiter plates were then incubated at 37 °C for 2 h and another 30 μL aliquot was removed from each well, serially diluted, and plated as described above. Plates were incubated overnight at 37 °C, and colony-forming units (CFUs) were enumerated the following day. Serum-mediated bacterial killing or growth was assessed by calculating the change in CFU over 2 h, normalized to the initial (0-h) count and the heat-inactivated serum control

### Animal husbandry

Female CD-1 mice at 29-31 g weight (5-7 weeks) were purchased from Charles River, UK. Upon arrival animals were independently randomised into high-efficiency particulate air (HEPA)–filtered cages in groups of 5. Sterile bedding, changed weekly, and enrichment were provided. Mice received food and water ad libitum and were housed in a 12hour/12hour light dark cycle. All mice were housed for an acclimatisation period of at least 1 week prior to undergoing any procedure. For each experiment, mice were randomly assigned to experimental groups. Investigators were not blind to the allocation.

### Intratracheal administration of inoculum and lung infection

The indicated strains were cultured overnight in LB to saturation and the inoculum was prepared in sterile PBS to the required concentration. Anaesthesia was induced by intraperitoneal injection of 80mg/kg ketamine and 0.8mg/kg medetomidine using a 27G 13mm needle (BD Microlance) and recovery after the intubation occurred at 32 °C following the administration of atipamezole (1 mg/kg). Pre-intubation body temperature was maintained by contact with a heat mat (Harvard Apparatus, U.K) and eye lubricant (2.0mg/g carbomer, Alcon UK) was applied. A 500 CFU/ 50 µl inoculum was prepared by dilution of saturated overnight cultures into sterile PBS. Intubation was achieved as previously described by placement of a 21G catheter (21G IV peripheral catheter (Insyte) BD Medical) using a fibreoptic intubation set (Kent Scientific) (Nat comms 2019).

An aliquot of inoculum was enumerated for each administration and all inoculums were ±10% of the stated figure. Mice recovered at 32°C (Warm Air System, Safety Cabinet Version red, VetTech, UK) until spontaneous movement and received 0.8mg/kg subcutaneous (BD Microlance 27G 13mm needle) atipamezole to reverse the ɑ-agonist into the neck scruff. After the mice resumed movement, they were returned to their individual ventilated home cages and monitored at regular intervals following infection. The experimental endpoint was 72 hours post-infection (hpi); mice that reached the human endpoint according to our approved animal protocol were humanely killed.

### Intraperitoneal infection

On the day of infection, the inoculum was prepared in sterile PBS from saturated overnights grown in LB at 150 CFU/200 µl. For the intraperitoneal injection, each mouse was gently restrained, and the lower right quadrant of the abdomen was injected with 200 µL of the inoculum at a shallow angle to avoid organ puncture, using a 27G, 13 mm needle. Mice were returned to their cages and closely monitored for signs of disease, in accordance with approved animal protocols. The experimental endpoint was 48 hpi; mice that reached the human endpoint according to our approved animal protocol were humanely killed.

### KP3 administration

The anti-MrkA human mAb KP3 and the control human IgG were administered 24 h prior to infection at a concentration of 15 mg/kg. Antibodies were administered via intraperitoneal injection using a 27G, 13 mm needle. None of the mice showed signs of distress or lost weight due to the mAb treatment alone.

### Cardiac puncture and blood processing

At the experimental end-point, animals were anesthetised via intraperitoneal injection of ketamine (100 mg/kg) and medetomidine (1 mg/kg) using 27G, 13 mm needles. Once anaesthesia was confirmed, blood was collected by cardiac puncture via a transdiaphragmatic inferior approach using 25G, 16 mm needles (BD Microlance). While still under anaesthesia, animals were humanely euthanized by cervical dislocation. A 20 µL aliquot of blood was then diluted into 180 µL of hypotonic lysis solution (1 mM EDTA in water) for bacterial enumeration and 2x 5 µL of blood were used for blood smears. Blood bacterial counts were determined by serial dilution in PBS followed by plating on LB agar containing rifampicin. Lung extraction or peritoneal lavage was performed as detailed in the corresponding sections; the rest of the blood was processed as detailed below depending on the aim of the experiment.

### For KP staining

The rest was transferred into a 50 mL centrifuge tube containing 5 mL (BALB/c) or 8 mL (CD-1) of red blood cell lysis buffer (0.15 M ammonium chloride, 0.01M potassium bicarbonate, 0.1 mM EDTA with 10 µM elinogrel), mixed well and incubated for 15 minutes at room temperature; the blood volume was recorded. PFA-PBS was then added (final concentration 1%) and samples were fixed on ice for 15 mins, after which 25 mL of cold PBS were added to each sample to dilute the PFA; samples were protected from light from here onwards. Blood samples were then centrifuged at 5000 xg for 10 mins, and pellets resuspended in DPBS, poured through a 70 µm strainer, washed once and resuspended in 0.4 mL DPBS.

### For determination of infection outcomes (Fig. 6)

The 25G needle was removed after blood collection to minimize haemolysis. The blood was then transferred into a serum collection tube containing a clot activator (BD Microtainer SST, #365968). Samples were left to clot at room temperature for 1 hour before centrifugation at 20,000 × *g* for 3 minutes. The resulting serum was aliquoted and stored at –80 °C until further analysis.

### Lung homogenate processing

Lungs were excised, weighed, and homogenized in gentleMACS C-tubes (Miltenyi Biotec) containing 3 mL of sterile PBS using a gentleMACS Octo Dissociator (Miltenyi Biotec) with the program *m_lung_2* run twice. An aliquot was separated for bacterial enumeration, determined by serial dilution in PBS followed by plating on LB agar containing rifampicin.

Triton X100 was then added to the rest to a final concentration of 0.5% and incubated at room temperature for 5 minutes to lyse all mammalian cells. PFA-PBS was then added to a final concentration 1% and samples were fixed on ice for 15 mins, after which they were transferred to a 50 mL centrifuge tube containing 25 mL of cold PBS; samples were protected from light from here onwards. Samples were centrifuged at 5000 xg for 10 mins and resuspended in tissue dissociation buffer (RPMI with 125 µg/mL liberase and 50 µg/mL Dnase I) and incubated with shaking at 37 °C for 30 mins in the dark. After incubation, EDTA was added to inactivate the enzymes and the samples were put through a 40 µm strainer, centrifuged, washed once in PBS and resuspended in 0.4 mL DPBS.

### Peritoneal lavage and downstream processing

Mice were secured supine on a dissection board using autoclave tape. 3 mL of DPBS with 3 mM EDTA and 2 mL of air were injected into the peritoneal cavity using a 25G (16 mm) needle to prevent leakage. After any procedures not affecting the peritoneum (e.g., BAL), the abdomen was opened to visualize fluid distribution. The board was tilted to ensure complete coverage. Peritoneal lavage fluid was collected using a 5 mL syringe fitted with an 18G (13 mm) needle inserted at an angle while the mouse was held upright. Fluid was withdrawn slowly without removing the needle, as leakage occurs at the puncture site. Fluid was transferred into a 15 ml falcon on ice and PFA was added to a final percentage of 1% to fix the samples for 15 mins on ice. Cold PBS was then added to a final volume of 15 mL to prevent over-fixation; samples were protected from light from here onwards. Lavage samples were then centrifuged at 5000 xg for 10 mins, and pellets resuspended in DPBS, washed once and resuspended in 0.2 mL DPBS.

### KP staining from *in vivo* samples and MrkA quantification

200 µl of blood KP samples, lung and peritoneal lavage KP samples in PBS processed as detailed above were transferred to 96 well V-bottom plates. Fixed overnights of sfGFP-expressing KP_WT_, KP_ΔmrkA_ and KP_mrkJ*_ were added as controls, in addition to extra samples from infected mice were included for unstained and fluorescence minus one controls for DAPI and MrkA (fluorescence minus one for KP expressing GFP provided by mock-infected samples).

Samples were centrifuged at 5000 *xg* for 5 mins and resuspended in 5% BSA-2mM EDTA supplemented with 10% normal donkey serum (NDS) in DPBS and incubated for 1 h to block non-specific finding. Samples were centrifuged, washed once in DPBS and resuspended in 2% BSA-DPBS with 2mM EDTA containing anti-MrkA antibody. Samples were then incubated overnight at 4°C on a microplate shaker (MTS 2/4 digital, IKA). The following day samples were washed two times in DPBS, resuspended in 2% BSA-2mM EDTA-DPBS containing anti-rabbit secondary and DAPI (to exclude any intact mammalian cells and stained debris) and incubated at room temperature shaking for 1 h. Samples were then centrifuged and washed two times with DPBS, resuspended in 200 µl 2% BSA-2mM EDTA-DPBS and kept in the dark until flow cytometry analysis. Flow cytometry analysis was performed on a Cytek Amnis CellStream after diluting the sample 1 in 5 for blood and lung; peritoneal lavage samples were not diluted. Positive (KP only) and negative (mock-infected) controls were used to gate on bacterial cells and a minimum of 1000 bacterial cells were counted per sample. Data analysis was carried out using FlowJo v10.10.0.

### Lung histology processing and immunostaining

The lower third of the left lung lobe was fixed in 4% paraformaldehyde (PFA) for 24 hours, then transferred to 70% ethanol. Fixed tissues were processed, embedded in paraffin, and sectioned at 5 μm thickness. Sections were stained with either haematoxylin and eosin (H&E) or prepared for immunofluorescence.

For immunofluorescence, sections were dewaxed sequentially in Histo-Clear (twice, 10 min each), 100% ethanol (twice, 10 min), 95% ethanol (twice, 10 min), 80% ethanol (twice, 3 min), and PBS with 0.1% Tween 20 and 0.1% saponin (PBS-TS; twice, 3 min). Antigen retrieval was performed by heating sections in demasking solution (0.3% trisodium citrate and 0.05% Tween 20, pH 6.0) for 30 minutes. After cooling, sections were blocked in PBS-TS with 10% NDS for at least 1 hour in a humidified chamber, followed by overnight incubation at 4 °C with primary antibodies diluted in the same blocking buffer. Primary antibodies are specified in the figure legends and in Table S4. Sections were then washed twice in PBS-TS (10 min each) and incubated with secondary antibodies (Table S4) and DAPI. After further washing, slides were mounted with ProLong Gold antifade reagent and cured overnight in the dark. Vector TruVIEW autofluorescence quencher was applied prior to mounting to minimize red blood cell autofluorescence. Images were captured using a Zeiss AxioVision Z1 microscope with an AxioCam MRm for H&E and a Hamamatsu camera for immunofluorescence. Image processing was performed with Zen 3.1 Blue software.

### Blood smear preparation and immunostaining

Blood smears were prepared on clean Superfrost microscope slides (VWR 631-0910) by placing 5 μL of anticoagulated blood in the centre of the coloured area. A plain push slide (Avantor VWR 631-1553) was used at a shallow angle to spread the drop using the push technique. Slides were air-dried and fixed in cold methanol for 10 minutes. Fixed slides were washed once with PBS, followed by a second wash with water to prevent PBS crystal formation, and then air-dried completely before staining or imaging.

A mini-PAP wax pen (ADI-950-232-0001, Enzo Life Sciences) was used to first delineate the area to be stained (usually middle of the smear)/ Fixed slides were washed 3 × 10 min in PBS. Non-specific binding was blocked with 300 µL blocking buffer (10% NDS, 2% BSA in PBS) for 45 min at room temperature in a humid chamber. Blocking buffer was removed without rinsing and primary antibodies were added diluted in 1% BSA/5% NDS/PBS and applied at 100 µL per slide overnight at 4°C in a humid chamber. Slides were washed twice with PBS, followed by incubation with fluorophore-conjugated secondary antibodies and DAPI diluted in 1% BSA/5% NDS/PBS (100 µL per slide) for 1 h at room temperature in the dark. After staining, slides were washed 3 × 10 min in PBS, then rinsed in water. Excess liquid was removed by blotting, and slides were mounted with Prolong anti-fade mounting medium and kept in the dark until imaging. The possibility of non-specific secondary binding of the secondary antibody was ruled out by staining using buffer without primary antibody to stain overnight. Samples were visualized using a Zeiss AxioVision Z1 microscope with a Hamamatsu camera and image processing was performed with Zen 3.1 Blue software

### Creatinine assay

Levels of the marker for kidney damage and decreased glomerular filtration rate, serum creatinine, were measured using the creatinine colorimetric assay from Abcam, following manufacturer’s instructions, and final measurements were expressed in mg/dL after transforming the interpolated values given as nmol/ 50 µl sample. Readings were acquired by measuring absorbance at 570 nm using a FLUOstar Omega plate reader. Serum was obtained as indicated above under the determination of infection outcomes section.

### Enzyme-linked immunosorbent assay (ELISA)

For analysis of serum cytokines, serum was obtained as indicated above under the determination of infection outcomes section. Serum samples were thawed on ice and serum cytokine levels were determined following the manufacturer’s instructions using the following: Mouse IL-6 uncoated ELISA kit, Mouse CXCL1/Kc DuoSet ELISA kit (Bio-Techne), or Mouse TNF uncoated ELISA kit.

### Data analysis and Visualization

No statistical methods were used to determine sample size. For ELISAs, two technical replicates were used to estimate experimental mean. Statistical analysis was performed by first testing for normality using the D’Agostino–Pearson test or the Kolmogorov–Smirnov test. If data were normally distributed, parametric tests were applied, using Welch’s t-test when variances were unequal. For non-normally distributed data, log-normality was assessed; if data followed a log-normal distribution, log-transformed parametric tests were used, again applying Welch’s correction when appropriate. If data were not log-normally distributed, non-parametric tests were used: Mann-Whitney for 2 groups, Kruskal-Wallis followed by Dunn’s post hoc test when 3 groups were compared. When more than three comparisons were made, p values were adjusted for multiple comparisons as indicated in the figure legends (Dunnett’s or Tukey’s post hoc test). Statistical analysis was only performed with datasets where each group analysed had two or more biological repeats. Data plotting and statistical analysis were performed using Prism 10.0.0 (GraphPad software Inc). Statistical details of experiments are described in the figure legends. A p value <0.05 is considered statistically significant. All schematics were made in Affinity Designer V2 (Serif Europe Ltd).

### Study approval

All animal studies were conducted at Imperial College London (Association for Assessment and Accreditation of Laboratory Animal Care accredited unit) in accordance with the Animals (Scientific Procedures) Act 1986 (Bleby 1989) under UK Home Office project license PP7392693. Ethical approval was obtained from the Imperial College Animal Welfare and Ethical Review Body. All procedures complied with relevant regulations for animal research, and results are reported in accordance with the ARRIVE guidelines (http://www.nc3Rs.org.uk/arrive-guidelines).

## Author contributions

**Julia Sanchez-Garrido:** Conceptualization, Data curation, Formal analysis, Investigation, Methodology, Supervision, Validation, Visualization, Writing – original draft, Writing – review & editing. **Sophia David:** Data curation, Formal analysis, Investigation, Methodology, Writing – review & editing. **Fabio Bagnoli:** Resources, Writing – review & editing. **Monia Bardelli:** Resources, Writing – review & editing. **Carlos Rodrigues dos Reis:** Resources, Writing – review & editing. **David M. Aanensen:** Funding acquisition. **Gad Frankel:** Conceptualization, Funding acquisition, Supervision, Writing – review & editing. **Joshua LC Wong:** Conceptualization, Data curation, Formal analysis, Investigation, Methodology, Supervision, Validation, Writing – review & editing.

## Acknowledgments

We thank the Flow Cytometry Facility at Imperial College London, in particular Dr Jessica Rowley and Dr Larissa Zárate García, for performing the FACS and always being so willing to help, and the histology facility for the histology processing. This study was supported by grants from The Wellcome Trust (224282/Z/21/Z) and the MRC (MR/R02671). S.D. and D.M.A. are supported by funding from the Centre for Genomic Pathogen Surveillance. The funders of the study had no role in study design, data collection, data analysis, data interpretation, or writing of the report.

## Data availability

Additional data are available in the supplementary material and from the corresponding author upon reasonable request.

